# Development and use of lentiviral vectors pseudotyped with influenza B haemagglutinins: application to vaccine immunogenicity, mAb potency and sero-surveillance studies

**DOI:** 10.1101/492785

**Authors:** Francesca Ferrara, George Carnell, Rebecca Kinsley, Eva Böttcher-Friebertshäuser, Stefan Pöhlmann, Simon Scott, Sasan Fereidouni, Davide Corti, Paul Kellam, Sarah C Gilbert, Nigel Temperton

## Abstract

Influenza B viruses cause respiratory disease epidemics in human populations and are included in seasonal influenza vaccines. Serological methods are employed to evaluate vaccine immunogenicity prior to licensure. However, the haemagglutination inhibition assay, which represents the gold standard for assessing the immunogenicity of influenza vaccines, has been shown to be relatively insensitive for the detection of antibodies against influenza B viruses. Furthermore, this assay, and the serial radial haemolysis assay are not able to detect stalk-directed cross-reactive antibodies. For these reasons there is a need to develop new assays that can overcome these limitations. The use of replication-defective viruses, such as lentiviral vectors pseudotyped with influenza A haemagglutinins, in microneutralization assays is a safe and sensitive alternative to study antibody responses elicited by natural infection or vaccination. We have produced Influenza B haemagglutinin-pseudotypes using plasmid-directed transfection. To activate influenza B haemagglutinin, we have explored the use of proteases by adding relevant encoding plasmids to the transfection mixture. When tested for their ability to transduce target cells, the newly produced influenza B pseudotypes exhibit tropism for different cell lines. Subsequently the pseudotypes were evaluated as surrogate antigens in microneutralization assays using reference sera, monoclonal antibodies, human sera collected during a vaccine immunogenicity study and surveillance sera from seals. The influenza B pseudotype virus neutralization assay was found to effectively detect neutralizing and cross-reactive responses despite lack of significant correlation with the haemagglutinin inhibition assay.

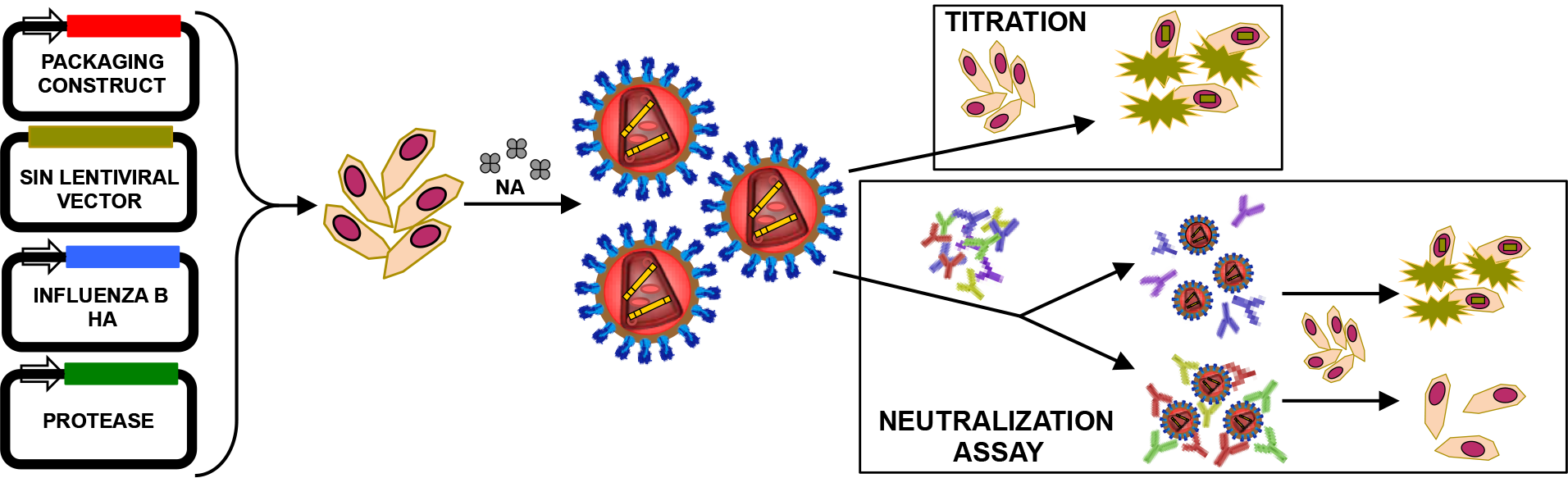

## Introduction

Every year respiratory disease epidemics cause severe illness and mortality in high-risk human populations, such as in children, the elderly, and individuals with underlying morbidity. Amongst the etiological pathogens causing these respiratory diseases, influenza viruses are one of the main contributors. Influenza viruses belong to the *Orthomyxoviridae* family and are enveloped negative sense single-stranded segmented RNA viruses. Three influenza virus *genera/types* (A, B, and C) able to infect humans can be distinguished; an additional influenza type (D) was recently proposed but it appears to be confined to animal populations (Hause et al. 2014). Influenza B has also been shown to infect seals (Osterhaus 2000). On the surface of these viruses are glycoproteins that have been used to characterise types and subtypes based on their antigenic properties. First, haemagglutinin (HA), is a homotrimeric type I glycoprotein, structurally characterized by a globular head that mediates viral attachment, and a stalk region. Neuraminidase (NA) is mushroom-shaped homotetramer that facilitates viral progeny release by cleaving cell-surface sialic acids. Influenza C and D encode a single glycoprotein type, the haemagglutinin-esterase-fusion (HEF), which is able to mediate both receptor-binding and receptor-destroying activities. The receptor of influenza A (H1-H16), B and C viruses, is protein-linked sialic acid. These are a family of carboxylated sugars that constitute the terminal monosaccharaides of animal protein glucidic residues (Varki 1992). If infection was not limited by other factors controlling viral replication, the HA would allow broad viral tropism, since sialic acids are present on almost all cell types regardless of species. These factors are mainly represented by restriction of the influenza polymerase complex activity, and by the activation of influenza HA. HA is synthesised as a single polypeptide precursor, HA0 (78kDa) that is then cleaved into two subunits HA1 (50-58 kDa) and HA2 (28-22 kDa). This HA cleavage, mediated by specific proteases, is necessary to expose a fusion peptide in the stalk region, and to permit HA conformational change after viral particle endocytosis. Subsequently, this permits envelope-endosome membrane fusion and the release of the viral core into the cell cytosol.

These factors however do not completely limit the influenza viruses’ ability to infect different animal species. For example, influenza A viruses are able to infect different avian and mammalian species; furthermore these viruses can cause human infections and have zoonotic and pandemic potential, since their antigenic characteristics are distinct from those of purely human influenza A viruses. Like influenza A, influenza B was reported to infect non-human mammalian species, such as seals, but these animal viruses are antigenically related to human viruses (Osterhaus 2000). For this reason, humans are considered the natural reservoir of influenza B and the zoonotic potential of influenza B animal viruses is considered limited. The fact that influenza B animal viruses have not shown zoonotic potential, and the lower infection and mortality rate associated with influenza B in comparison to influenza A has resulted in influenza B being understudied.

The clinical and public health community has until recently underestimated the influenza B burden. It has become evident that it is necessary to raise awareness of influenza B virus infection, especially by increasing epidemiological surveillance and seasonal vaccination coverage (Glezen et al. 2013). Influenza B strains are included in seasonal vaccines, but frequent mismatches between vaccine and circulating strains has left the at-risk population unprotected (Dolin 2013). This is partially due to the fact that in the late 1970s, a large-scale genomic reassortment event involving all eight influenza B segments led to the generation of two distinct influenza B lineages: the Victoria lineage (prototype virus B/Victoria/2/1987) and the Yamagata lineage (prototype virus B/Yamagata/16/1988) (Chen & Holmes 2008; Rota et al. 1990). These two lineages continue to diverge, are subject to reassortment, and to co-circulation, even if one tends to dominate over the other for a determinate period of time (Chen & Holmes 2008). To resolve the vaccine mismatch issue, quadrivalent vaccines containing one representative strain for each influenza B lineage together with the influenza A H1N1 and H3N2 viruses were developed and were recently licensed (Tinoco et al. 2014; Beran et al. 2013; Pépin et al. 2013). Despite the improved vaccine coverage, epidemiological surveillance and rigorous vaccine testing are still needed (Eichner et al. 2014; Beran et al. 2013). Classical serological assays, such as haemagglutination inhibition (HI), Single Radial Haemolysis (SRH), and microneutralization (MN), are usually employed for this purpose. Nevertheless, several studies have independently shown that the HI assay is insensitive for the measurement of Influenza B seroconversion, since it routinely underestimates antibody titres in comparison to SRH (Mancini et al. 1983; Oxford et al. 1982; Wood et al. 1994). Ether-treatment of the antigen (virus) during HI was shown to increase sensitivity (Pyhälä et al. 1985; Kendal & Cate 1983; Monto & Maassab 1981), however this increase is insufficient to attain SRH and MN consistency (Ansaldi et al. 2004; Kendal & Cate 1983; Mancini et al. 1983).

Additionally, classical serological methods do not enable the efficient detection of HA stalk-directed antibodies (Dreyfus et al. 2013; Corti et al. 2011; Wang et al. 2010; Ekiert et al. 2012; Ekiert et al. 2009; Sui et al. 2009; Okuno et al. 1993) that are promising targets for influenza ‘universal’ vaccines, a frequently discussed and reviewed subject (de Vries et al. 2015b; de Vries et al. 2015a; Krammer & Palese 2015). With the evidence that antibodies able to neutralize the two influenza B lineages and in some cases, also influenza A viruses are detectable in humans (Yasugi et al. 2013; Dreyfus et al. 2013), interest in studying the cross-neutralizing response from an influenza B perspective has increased. Monoclonal antibodies can be employed to delineate these responses, whereupon lineage specific or broader epitopes can be identified and tested, yielding data which wouldn’t be possible using polyclonal sera.

In recent years the use of retroviral vectors (including lentiviral vectors based on the HIV-1 genome) pseudotyped with the HAs of influenza A viruses has been shown to be effective for studying antibody responses especially against high-containment viruses with pandemic potential (Carnell et al. 2015). Influenza A HA pseudotypes (PV) are useful tools to study influenza heterosubtypic antibody responses, since it is thought that a lower density of the HAs on the vector surface permits increased access to HA-stalk directed antibodies, the major mediator of the heterosubtypic antibody response (Corti et al. 2011). With this knowledge, and to address the aforementioned serological assay issues, we have developed a panel of lentiviral vectors pseudotyped with influenza B HAs. These newly established influenza B PV reagents, which have hitherto not been reported in the literature, were subsequently investigated for their feasibility of use as surrogate antigens in neutralization assays, and to study cross-neutralizing antibody responses. A prerequisite for the production of high titre pseudotypes requires understanding the life cycle of the virus that will donate its envelope protein. HA activation through proteolytic cleavage is essential in the influenza virus life cycle since it permits low-pH dependent fusion with the endosome-membrane after attachment and endocytosis of the virus, allowing the release of the viral core into the cytosol. For influenza B, HA activation has been observed in embryonated chicken eggs (Zhirnov et al. 1994), and in a trypsin-dependent and independent manner in MDCK cells (Noma et al. 1998; Lugovtsev et al. 2013). Additionally, experiments have shown that the NA could also have a role in HA activation by removing glucidic residues on the HA surface (Yamamoto-Goshima & Maeno 1994; Shibata et al. 1993). Furthermore, it was shown that influenza B HA can be cleaved by porcine pancreatic elastase when the cleavage-linked arginine is substituted with an alanine or a valine (Stech et al. 2011). Influenza B HA can also partially support a poly-basic cleavage site that permits the cleavage by subtilisin-like proteases (Brassard & Lamb 1997). More recently, HAT and TMPRSS2 have been shown to be able to cleave and activate influenza B HA in *in vitro* models (Böttcher-Friebertshäuser et al. 2012). Data on further proteases that could be involved in the cleavage of influenza B HA in nature are lacking.

## MATERIALS AND METHODS

### Cell lines

The Human Embryonic Kidney (HEK) 293T/17 cells were used for PV production and as target cell lines. They were maintained in Dulbecco’s Modified Eagle Medium (DMEM, PAN-Biotech) supplemented with 15% (v/v) Foetal Bovine Serum (FBS, PAN-Biotech) and 1% (v/v) penicillin-streptomycin (Sigma-Aldrich).

Madin-Darby Canine Kidney (MDCK) and A549 cells were additionally used as target cell lines to study influenza B PV transduction and tropism. Cells were cultured in DMEM supplemented with 5% FBS and 1% of penicillin-streptomycin and in DMEM/F12 media (Hyclone) with 10% FBS and 1% penicillin-streptomycin, respectively.

### HA-encoding, protease-encoding, packaging, and lentiviral vector plasmids

B/Bangladesh/3333/2007 and B/Brisbane/60/2008 HA genes were amplified from cDNA and subcloned into plasmid expression vector pI.18 (Temperton et al. 2007; Cox et al. 2002). B/Hong Kong/8/1973, B/Victoria/2/1987, B/Yamagata/16/1988 and B/Florida/4/2006 codon optimized HA genes were synthesized by GenScript and cloned into phCMV1 as previously described (Corti et al. 2011). pCAGGS-HAT (Human airway trypsin-like protease), pCAGGS-TMPRSS2 (type II transmembrane protease serine 2, Böttcher et al. 2006), and pCMV-TMPRSS4 (Bertram et al. 2010) expressing respectively the human airway trypsin-like protease (HAT), (TMPRSS2), and type II transmembrane protease serine 4 (TMPRSS4) were used to activate the influenza B HA during lentiviral vector production. The second-generation packaging construct pCMVΔR8.91, and the self inactivating lentiviral vectors pCSFLW expressing firefly luciferase and pCSemGW (kindly provided by Greg Towers, University College London, UK) expressing emerald green fluorescence protein (emGFP) were used for PV production.

### Production of influenza B PVs

Influenza B PVs were produced in 6-well plates using branched polyethylenimine co-transfection of 500 ng of packaging construct, 750 ng of lentiviral vector, and 500 ng of HA-encoding plasmid. Additionally, 125 ng or 250 ng of protease-encoding plasmid were added to the DNA transfection mix to allow influenza B HA activation and cleavage. As controls, influenza PVs were additionally produced in the absence of protease-encoding plasmid (Δ protease). 48 hours post-transfection, PVs were collected, filtered using 0.45 μm mixed cellulose ester membrane filters (Merck Millipore), and titrated.

### Titration of influenza B PVs and transduction of different cell lines

Titration experiments of firefly luciferase-expressing or emerald GFP PVs were performed in Nunc™ F96 MicroWell™ white or transparent polystyrene plates (Thermo Fisher Scientific). The PV production titre was evaluated by transducing HEK293T/17 cells with 2-fold serial dilutions of PV, performed in a total mix volume of 100 μl, discarding 50 μl from the last dilution. Then 1.5×10^4^ HEK293T/17 cells were added to each well of the 96-well titration plate. As a control for HA cleavage, Δ protease influenza B PVs were also activated by addition of 1 mg/ml (TPCK)-trypsin (Sigma-Aldrich) to each 2-fold-dilution well, to a final concentration of 100 μg/ml. The plate was incubated at 37°C 5% CO_2_, and after 30 minutes 50 μl of trypsin neutralizing solution (Lonza) was added to each well. Subsequently, the HEK293T/17 cell suspension was also added to the plates. After 48 hour incubation at 37°C 5% CO_2_, firefly luciferase gene expression was evaluated and quantified by luminometry using the Bright-Glo™ assay system (Promega) and GloMax Multi detection system luminometer (Promega). Each relative luminescence unit (RLU) value obtained at different PV dilution points was transformed into RLU/ml; the arithmetic mean of these concentrations was considered as the PV production titre (expressed as RLU/ml). Influenza B PVs produced after activation/cleavage-optimization and expressing firefly luciferase or emerald GFP were also tested, using an equivalent protocol, for their ability to transduce two further target cell lines: MDCK and A549.

### Serum samples and monoclonal antibodies

NIBSC 11/136 anti-B/Brisbane/60/2008 serum (NIBSC) was used as the primary antibody in western blot experiments and as positive control in neutralization assays. Sera obtained during the clinical trial NCT00942071 (Antrobus et al. 2013) were used in PV neutralization assays (pMN). These sera were collected from adults aged 50 years and over pre- and post- administration of an influenza split virion vaccine (Sanofi Pasteur MSD, France) containing HAs of A/California/7/2009 (H1N1), A/Perth/16/2009 (H3N2), and B/Brisbane/60/2008. Nine subjects received an injection of Modified Vaccinia Ankara (MVA) vectors expressing the A/Panama/2007/1999 (H3N2) NP and M1 (MVA-NP+M1) antigens as a single fusion protein (Antrobus et al. 2013; Berthoud et al. 2011). The remaining subjects (n=8) received a TIV and saline placebo. These sera were previously analysed (Antrobus et al. 2013) in an HI assay against B/Brisbane/60/2008 and the data obtained were employed for a correlation study with B/Brisbane/60/2008 PV neutralization assay. The seal serum samples (n=8) utilized were taken from seals caught in the Caspian Sea (CS) from 2006-2010, at the Europe/Asia border. Neutralisation of the full set of influenza B PV was evaluated using a panel of 5 previously characterised monoclonal antibodies from the Division of Viral Products, Food and Drug Administration, USA, see Table 1 (Verma 2017).

**Table 1.** Influenza-B lineage specific monoclonal antibodies (Verma 2017)

|  |  |  |  |  |  |
| --- | --- | --- | --- | --- | --- |
| Specificity | Victoria |  | Yamagata |  | Both Lineages |
| Designation | 8E12 | 5A1 | 3E8 | 1H4 | 2F11 |

### Western blotting

To study the activity of different proteases on the HAs of the influenza B PVs and to confirm HA cleavage, a western blot was performed using B/Brisbane/60/2008. Briefly, 2 ml of PV were centrifuged on a fixed-angle centrifuge at 3000 g and 4°C for 24 h. Then supernatant was removed and 150 μl of OptiMEM^®^ I reduced serum medium was added to the 2 ml tubes (Thermo Fisher Scientific, cat. n° 11519934) that were incubated overnight at 4°C to permit PV resuspension. After this, PVs were stored at −80°C before preparing the samples for sodium dodecyl sulphate-polyacrylamide gel electrophoresis (SDS-PAGE). As HA activation/cleavage control, TPCK-Trypsin treatment on the Δ protease influenza B PVs was accomplished using TPCK-Trypsin to a final concentration of 100 μg/ml.

A 4-15% Mini-PROTEAN^®^ TGX™ precast polyacrylamide gel (Bio-Rad Laboratories) was used for the SDS-PAGE of reduced and denatured influenza B PVs following the manufacturer’s instructions. The gel was transferred onto an Immuno-Blot^^®^^ low fluorescence polyvinylidene fluoride membrane (PVDF, Bio-Rad Laboratories) for 60 min at 100 V. The membrane was blocked overnight using 10% (w/v) dried milk + 0.01% (v/v) Tween20-PBS. The anti-B/Brisbane/60/2008 serum (NIBSC 11/136) was used as the primary antibody, diluted 1:500 in 5% (w/v) dried milk-0.01% (v/v) Tween20-PBS. The membrane was washed abundantly with 0.01% (v/v) Tween20-PBS prior to addition of donkey antibody (anti-sheep/goat). IgG Dylight^®^800 antibody diluted 1:20000 5% (w/v) dried milk-0.01% (v/v) Tween20-PBS was added to detect the sheep-origin antisera. After washing to remove unbound secondary antibody, the PVDF membrane was detected using the Odyssey^®^ Sa Infrared Imaging System (LI-COR Bioscience) at 800 nm.

### Pseudotype based microneutralization assay

The neutralization activity of the positive control NIBSC 11/136 anti-B/Brisbane/60/2008 serum and mAbs were evaluated against three different influenza B PVs (B/Brisbane/60/2008, B/Hong Kong/8/1973, and B/Florida/4/2006) by performing pMN in quadruplicate according to established protocol (Ferrara & Temperton 2018). Briefly, 2-fold serial dilutions of serum were performed in a Nunc™ F96 MicroWell™ white polystyrene plate. The plate was centrifuged for 1 min at 500 g using an ELMI CM-6MT Centrifuge and rotor 6M04, subsequently 50 μl of influenza B PV solution with a concentration 1×10^6^ RLU/50 μl was added to each well. The plate was centrifuged for 1 min at 500 g before incubation at 37°C 5% CO_2_. After 1 hour incubation, 1.5×10^4^ HEK293T/17 cells were added to each well and, after a final 1 min centrifugation at 500 g, the plate was incubated at 37°C 5% CO_2_. 48 h later firefly luciferase gene expression was evaluated. To determine if the pMN assay correlates with standard serological assays (e.g. HI), to investigate the suitability of influenza B pMN assays in a vaccine immunogenicity study, and to investigate if it is possible to detect cross-reactive responses to influenza B, the human sera from clinical trial NCT00942071 were also tested in pMN assay. Additionally, the neutralization activity of a panel of seal sera was evaluated (table 2). A 1:100 starting dilution of seal sera was used for neutralization against B/HK, B/Yam and B/Vic. Positive control from NIBSC 11/136 sheep, anti-B/Brisbane/60/2008 serum (starting dilution 1:1000). Human positive controls were obtained from Emanuele Montomoli at the University of Siena (starting dilution 1:1000). The negative control used was FBS (starting dilution 1:100). For all assays, PV input was 1 × 10^6^ RLU/well.

**Table 2.**
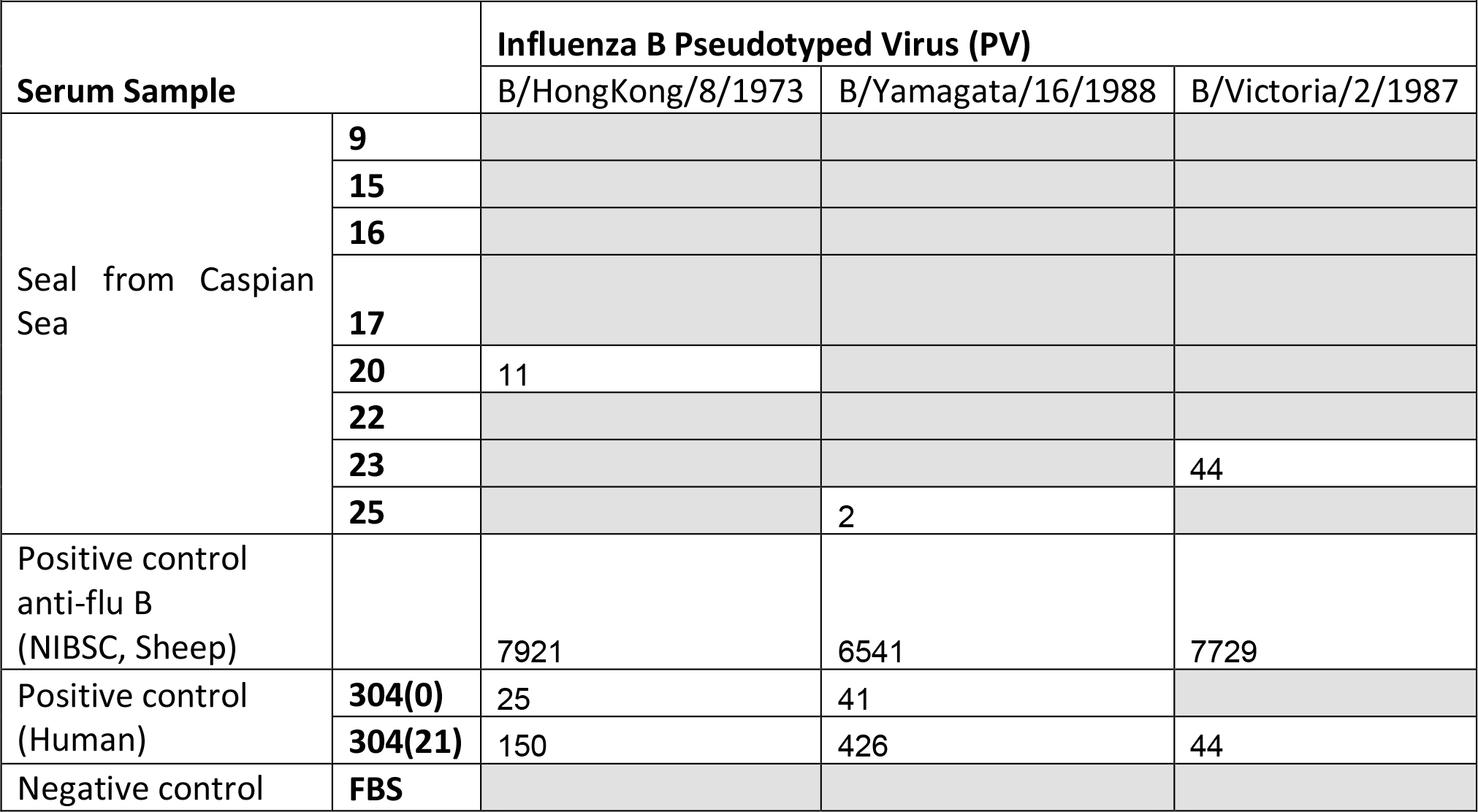
Use of influenza B PV in neutralization assays utilizing seal sera. Values reported as reciprocal dilution values for serum to neutralize 50% of input PV (IC_50_). Grey cells indicate no neutralization

|  |  |  |  |  |
| --- | --- | --- | --- | --- |
| Serum Sample |  | Influenza B Pseudotyped Virus (PV) |  |  |
|  |  | B/HongKong/8/1973 | B/Yamagata/16/1988 | B/Victoria/2/1987 |
| Seal from Caspian Sea | 9 |  |  |  |
|  | 15 |  |  |  |
|  | 16 |  |  |  |
|  | 17 |  |  |  |
|  | 20 | 11 |  |  |
|  | 22 |  |  |  |
|  | 23 |  |  | 44 |
|  | 25 |  | 2 |  |
| Positive control anti-flu B (NIBSC, Sheep) |  | 7921 | 6541 | 7729 |
| Positive control (Human) | 304(0) | 25 | 41 |  |
|  | 304(21) | 150 | 426 | 44 |
| Negative control | FBS |  |  |  |

### Statistical analysis

Data analysis of pMN results was performed using Microsoft^®^ Excel 2011 and GraphPad Prism^®^ version 6 (GraphPad Software). To measure neutralization activity, RLU results were normalized and expressed as the percentage of neutralization (inhibition of PV entry into cells as indicated by reduction in luminescence) using the arithmetic mean of the RLU values of viral input control and of the cell control as 0% and 100% neutralization values respectively. The half maximal inhibitory concentration (IC_50_) was calculated using a non-linear regression method (GraphPad Prism^®^) and was expressed as the reciprocal dilution factor in which the sample shows 50% neutralization activity.

For the NCT00942071 trial, Pearson correlation between the log_10_ HI assay titre and the log_10_ IC_50_ antibody titre values was performed using GraphPad Prism^®^. IC_50_ titres of the NCT00942071 sera were reported in Box-and-Whisker plots to allow a graphical comparison of the results. A non-parametric Wilcoxon matched-pairs signed rank test to assess statistical significance between day 0 and day 21 in the clinical trial was applied considering that the data were not following Gaussian distributions. Clinical trial data were also then stratified on the basis of the vaccine regimen: TIV + placebo or TIV + MVA-NP+M1, to evaluate the effect of the MVA-NP+M1 in broadening the heterosubtypic antibody response. To avoid multiple comparisons, statistical analysis was performed using the IC_50_ fold-increase between TIV + placebo and the TIV + MVA-NP+M1 groups in a Mann-Whitney U test.

### Bioinformatic analysis

Influenza B HA gene sequences that were used in PV production were used to calculate percentage identity between amino acid sequences, and for phylogenetic analysis. First, HA-encoding sequences were downloaded from the Influenza Virus Resource database (IVRD). Then, percentage identity between amino acid sequences was calculated by pair-wise alignment using Jalview (Waterhouse et al. 2009).

For phylogenetic analysis, codon-based alignment was performed on the sequences using the MUSCLE algorithm (Edgar 2004) in Molecular Evolutionary Genetics Analysis (MEGA) (Tamura et al. 2011). Bayesian evolutionary analysis by sampling trees (BEAST) software package was used to build the phylogenetic tree (Drummond & Rambaut 2007; Drummond et al. 2012). The Hasegawa, Kishino and Yano (HKY) + Gamma model, which was determined as the best fit model by Jmodetest analysis, was used as nucleotide substitution model; furthermore the year of strain isolation was added as a parameter to permit the software to evaluate the time-dependent rates of molecular evolution (molecular clock), calculate branch length and incorporate a time-scale in the tree. The maximum clade credibility tree was calculated by discarding (burn-in) the first 25% of saved trees. The tree generated was then graphically elaborated with FigTree (http://tree.bio.ed.ac.uk/software/figtree/) and with MEGA.

## RESULTS AND DISCUSSION

### Co-transfection of a protease-encoding plasmid is necessary for production of high titre influenza B PVs

In this study, we show that for production of high titre influenza B PVs it is necessary to activate the influenza B HA using a protease. Here we have analysed three different proteases that are associated with influenza A HA cleavage and that were previously reported to be necessary to produce high titre influenza A PVs: HAT, TMPRSS2, and TMPRSS4 (Ferrara et al. 2012; Bertram et al. 2010). Figure 1 shows that transduction titres of different influenza B PVs are dependent on the type of proteases used and on the quantity of protease-encoding plasmid co-transfected with a fixed amount of the HA-encoding plasmid, the packaging construct, and the lentiviral vector construct.

**Figure 1.**
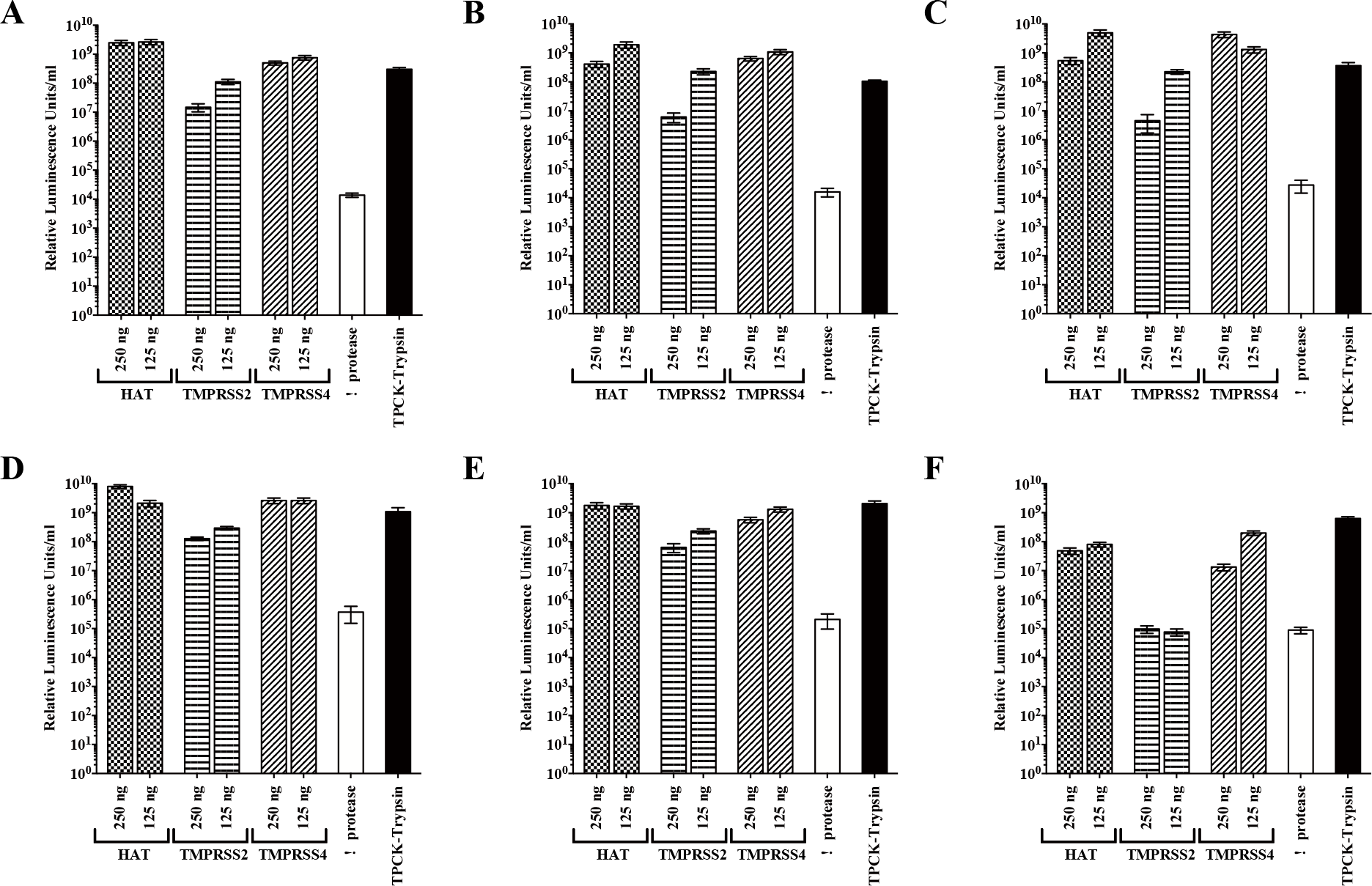
Production of influenza B PV. Different quantities (250 ng and 125 ng) and types (HAT, TMPRSS2, and TMPRSS3) of protease-encoding plasmids were co-transfected with packaging constructs, lentiviral vectors, and plasmids expressing the HA of B/Hong Kong/8/1973 (A) B/Victoria/2/1987 (B), B/Yamagata/16/1988 (C), B/Florida/4/2006 (D), B/Bangladesh/3333/2007 (E), and B/Brisbane/60/2008 pseudotypes (F). Pseudotypes produced in absence of protease-encoding plasmids and subsequently activated using TPCK-Trypsin are also shown. Production titres expressed as RLU/ml.

In the absence of protease, influenza PVs are produced, but only a small proportion of HA is cleaved by endogenous proteases, leading to low overall transduction competence. In contrast, when the HAT-encoding plasmid is added to the transfection mix, the PV HAs are activated to a greater degree. In the presence of this protease, PV transduction titre reached 10^9^ RLU/ml. The TMPRSS2-activated influenza B PVs have lower transduction titres, which are still approximately a log lower (bar B/Brisbane/60/2008) but still significantly above Δprotease values. The role of TMPRSS4 in influenza B HA cleavage/activation is also shown: the transfection of the encoding plasmid during influenza B PV production is associated with high titres.

### HAT, TMPRSS2, TMPRSS4 and TPCK-Trypsin activate influenza B PVs via HA cleavage

To confirm that the addition of a protease-encoding plasmid mediates HA cleavage, a western blot was performed on B/Brisbane/60/2008 PVs (Figure 2). In all the B/Brisbane/60/2008 PVs produced with protease-encoding plasmid addition, bands corresponding to HA1 can be observed in the western blot. HAT and TMPRSS4 produced PVs displaying a wider HA1 band, whereas the bands observed when TMPRSS2 is used are thinner. The band intensities correlate with the PV titres in which TMPRSS2 viruses show lower titres compared to HAT and TMPRSS4 produced PVs. Furthermore, in the western blot, the a protease PV shows not only a band at ~78 kDa corresponding to HA0, but also a small band at ~55 kDa. The presence of the HA1 band is likely to be related to the presence of proteases in the producer HEK293T/17 cells and explains the entry of a protease PV into HEK293T/17 target cell line observed during titration experiments. The western blot shows that the HA cleavage mediated by proteases can also be reproduced through TPCK-trypsin treatment of the Δ protease PVs: the HA0 band was replaced by bands corresponding to HA1 and HA2. These results clearly support the hypothesis that the three proteases and TPCK-trypsin cleave the HA0 into two subunits HA1 and HA2.

**Figure 2.**
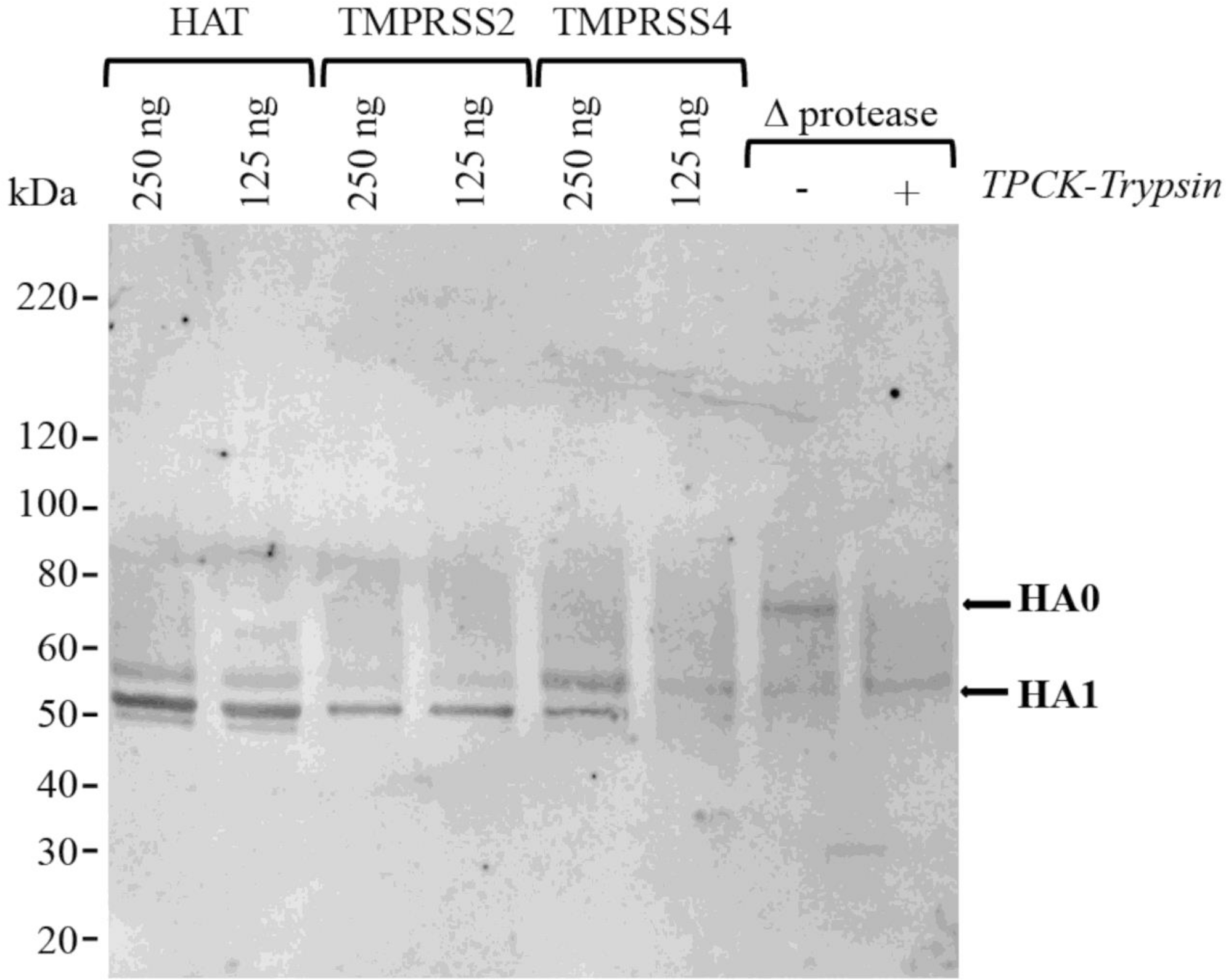
Cleavage of influenza B HA PV using HAT, TMPRSS2/TMPRSS4 and TPCK-Trypsin. Western blot analysis reveals differences in glycosylation profiles for the products of HA0 cleavage: HA1, HA2.

Interestingly, in the western blot, the HAs activated by HAT, TMPRSS2 and TMPRSS4 present heterogeneous glycosylation characteristics: more than one band between 50 kDa and 60 kDa, all corresponding to HA1, are present. Instead, in the Δ protease pp before and after TPCK-trypsin treatment only a single band at 55 kDa is observed. Elsewhere it was noted that exogenous bacterial NA can play a role in removing glucidic residues from the HA surface, to enhance viral attachment to the receptor(Brassard & Lamb 1997). However, this it is not the case since Δ protease pp do not show heterogeneous glycosylation characteristics and were also treated with exogenous bacterial NA during PV production. For this reason, the most likely hypothesis is that the proteases (especially TMPRSS2) that mediate HA cleavage intracellularly (Böttcher et al. 2009), could interact with the cell glycosylation machinery (Walker et al. 1992).

### Influenza B PVs enter into different target cell lines

The influenza B PV produced were also tested for their ability to transduce HEK293T/17, MDCK, and A549 cells. In the quantitative results obtained using firefly luciferase-expressing lentiviral vector particles (Figure 3), all the influenza B PV tested can transduce the target cell lines examined: higher transduction titres were obtained when the producer cells HEK293T/17 were used, whereas the classical influenza producing cell lines MDCK and the human lung carcinoma A549 show lower transduction titres. These results were also confirmed using emGFP-expressing influenza B PV (Figure 4), demonstrating that the influenza B PV can transduce HEK293T/17, MDCK, and A549 cells.

**Figure 3.**
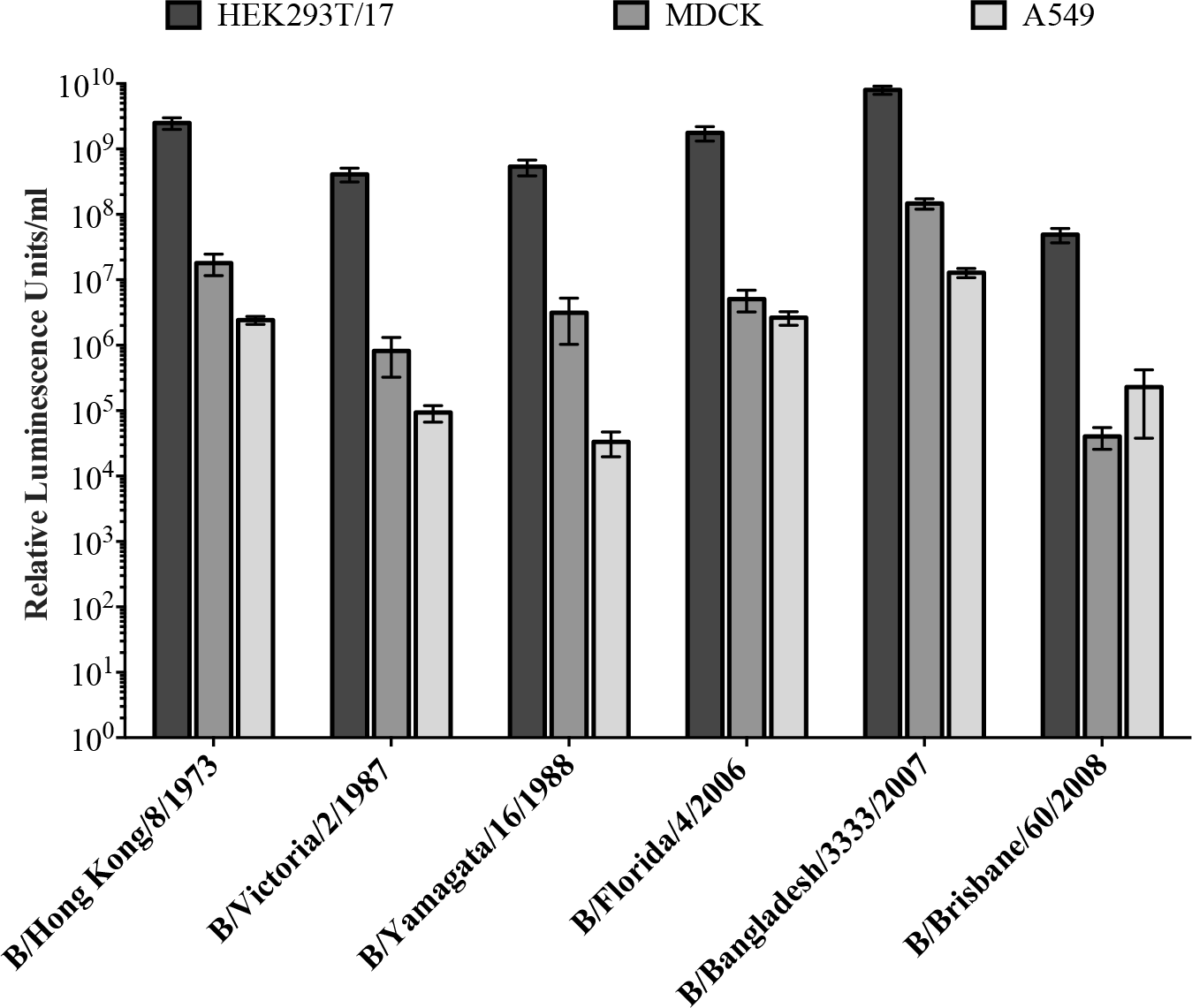
Influenza B luciferase-expressing pseudotypes transduce different cell lines. Multiple cell lines can be transduced by influenza B bearing lentiviral pseudotypes, with optimal luciferase production in HEK293T/17 cells, followed by MDCK and A549.

**Figure 4.**
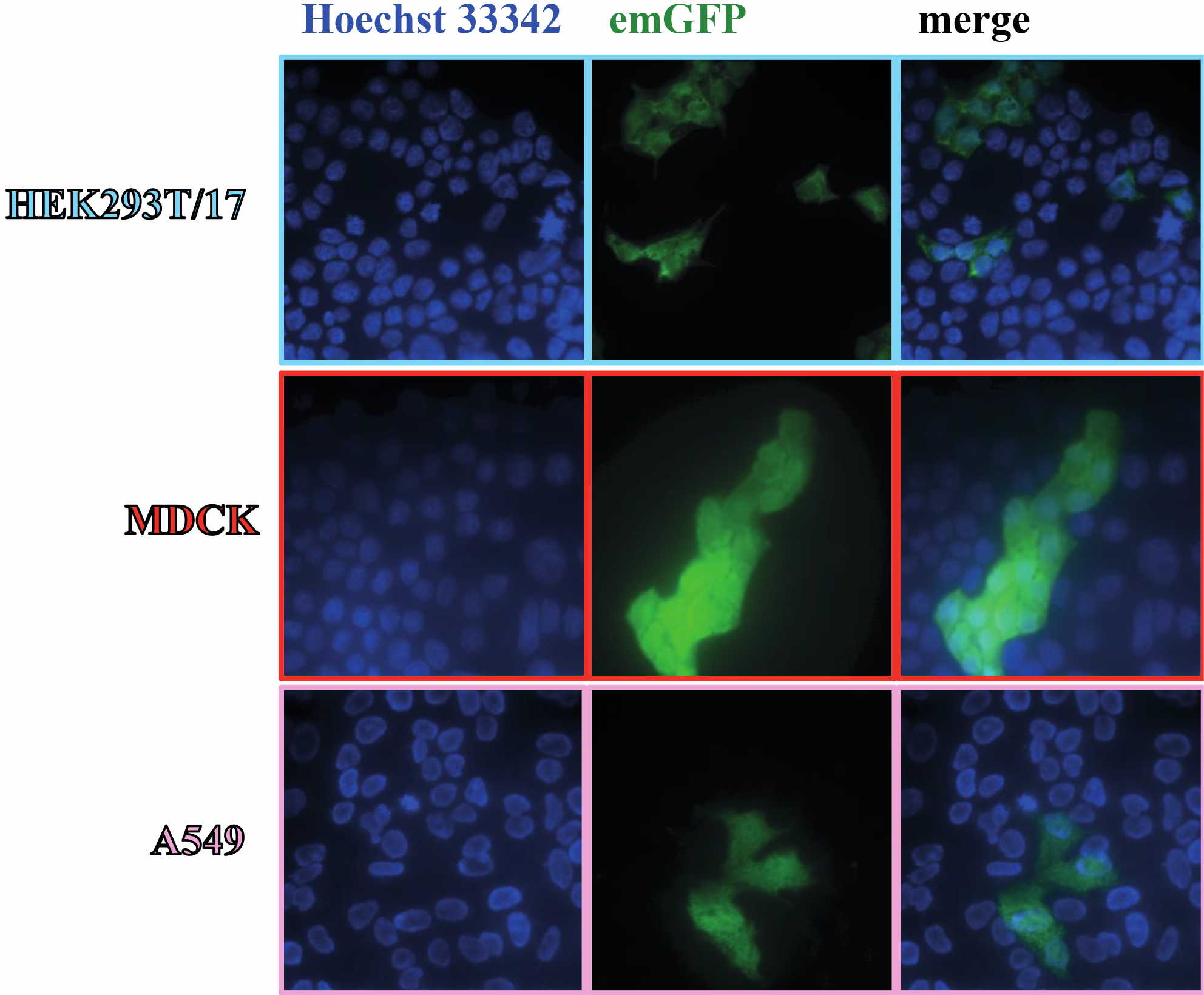
B/Florida/4/2006 emGFP-expressing pseudotypes transduce different cel lines (HEK293, MDCK and A549) Influenza B lentiviral pseudotypes (strain B/Florida/4/2006) containing an emerald GFP reporter successfully transduced HEK293T/17, MDCK and A549 cells

There are different factors that could explain why different HAs show better entry capacity than others in different cell lines. Firstly, influenza B HA recognises different sialoglycan residues present on the glycoprotein surface to mediate cell entry, and different strains/viruses can have different preferences (Wang et al. 2012; Xu et al. 1994; Matrosovich et al. 1993); secondly cells can present a different proportion of sialic acid combinations (Lugovtsev et al. 2013; Varki & Varki 2007; Svennevig et al. 1995). This could be important for entry into certain cell lines, in relation to differential sialic acid distribution between cells (Sieben et al. 2012). However, these elements alone cannot explain the differences in titre observed, especially taking into consideration that MDCK should be a highly susceptible cell line since it is routinely used for influenza virus infection and amplification.

Here, HEK293T/17 are used as producer cells and subsequently as target cells, exhibiting higher transduction efficiency compared to the other cell lines tested. Pizzato *et al* and Voelkel *et al* have observed that lentiviral vector particles are able to bind to the cell membrane and be subjected to endocytosis in absence of envelope proteins (Pizzato et al. 1999; Voelkel et al. 2012). In the cases reported as here, the producer and target cell lines were of the same origin; furthermore, this nonspecific binding was related especially to human cell lines. It is reasonable to enquire if the identical origin of the PV lipid bilayer and of the target cell plasma membrane can influence the transduction activity. The lipid bilayer and the characteristics of the membrane lipids can play an important role in PV entry as lipid properties have been shown to play an important role in membrane fusion (Heaton & Randall 2011; Sun & Whittaker 2003; Chernomordik et al. 1998).

### Influenza B PV are neutralized by reference antiserum and lineage specific monoclonal antibodies

The use of influenza B PVs as surrogate antigens in neutralization assays was investigated. First, the ability of a reference antiserum to neutralize three of the six PVs generated: B/Brisbane/60/2008, B/Florida/4/2006, and B/Hong Kong/8/1973, was evaluated. B/Brisbane/60/2008 and B/Florida/4/2006 PVs were chosen as they represent strains of co-circulating influenza B lineages previously or currently used in influenza seasonal vaccination. B/Hong Kong/8/1973 PV was chosen as a representative strain circulating before the influenza B lineage division (Figure 5). The reference serum used neutralized the PVs tested (Figure 6). It was also able to differentiate between the different PVs, showing correlation with amino acid identity and different neutralization activities. The serum neutralizes the homologous B/Brisbane/60/2008 most efficiently, followed by B/Hong Kong/8/1973, which shares 94.9% amino acid identity with B/Brisbane/60/2008, and B/Florida/4/2006 (93.5% amino acid identity). This could indicate that, as already observed with influenza A PVs (Molesti et al. 2014; Valentich 1981; Corti et al. 2010; Corti et al. 2011; Garcia et al. 2009), the pMN assay is a sensitive assay, detecting cross-neutralizing antibody responses, while maintaining specificity.

**Figure 5.**
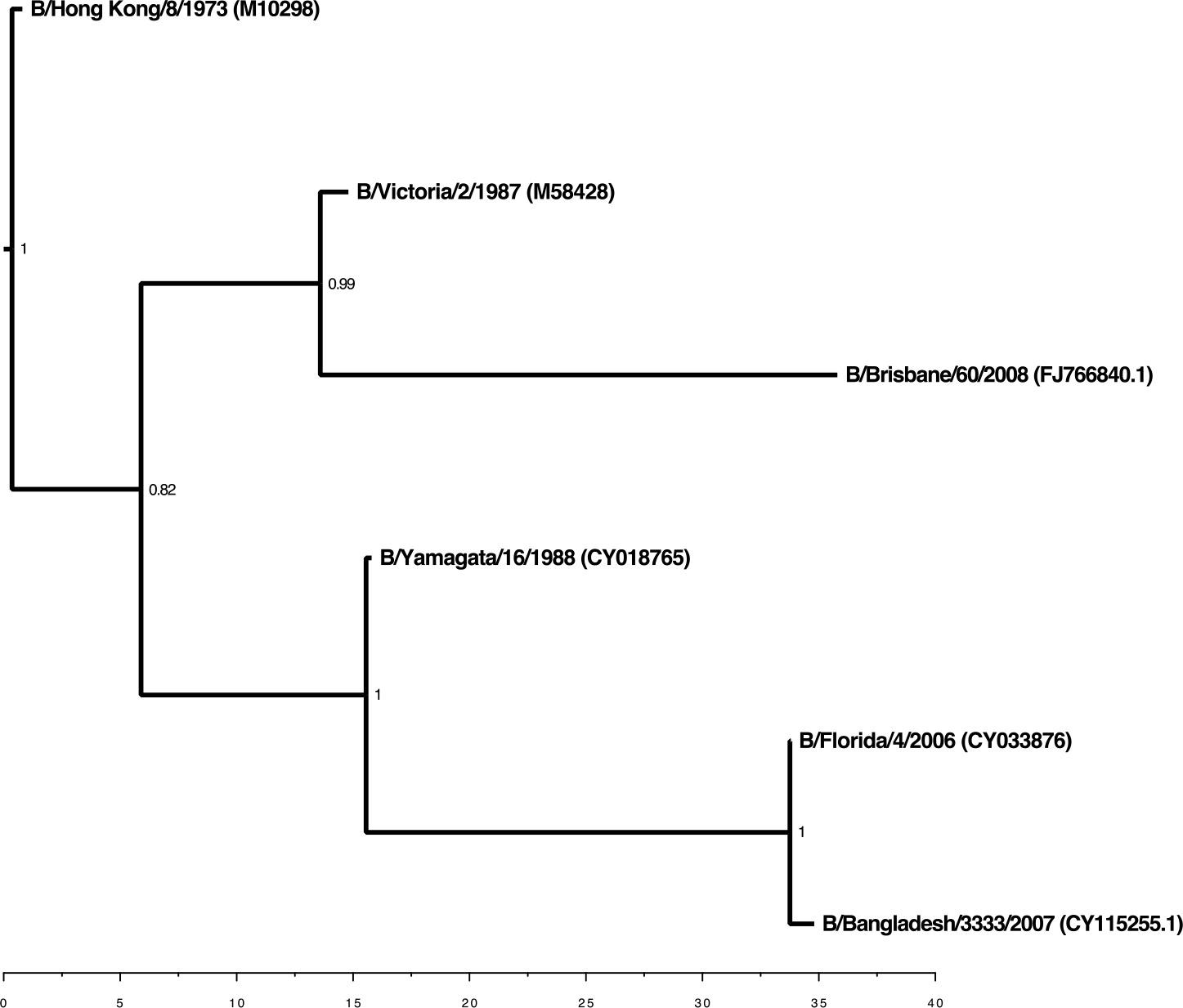
Phylogenetic tree of the influenza B HAs used for pseudotype production. This tree shows the Victoria lineage and Yamagata lineage division. Accession numbers are reported with the strain name on the tree tips. Posterior probabilities are reported on the nodes. Axes represent time scales with origin at the most recent circulating strain (B/Brisbane/60/2008).

**Figure 6.**
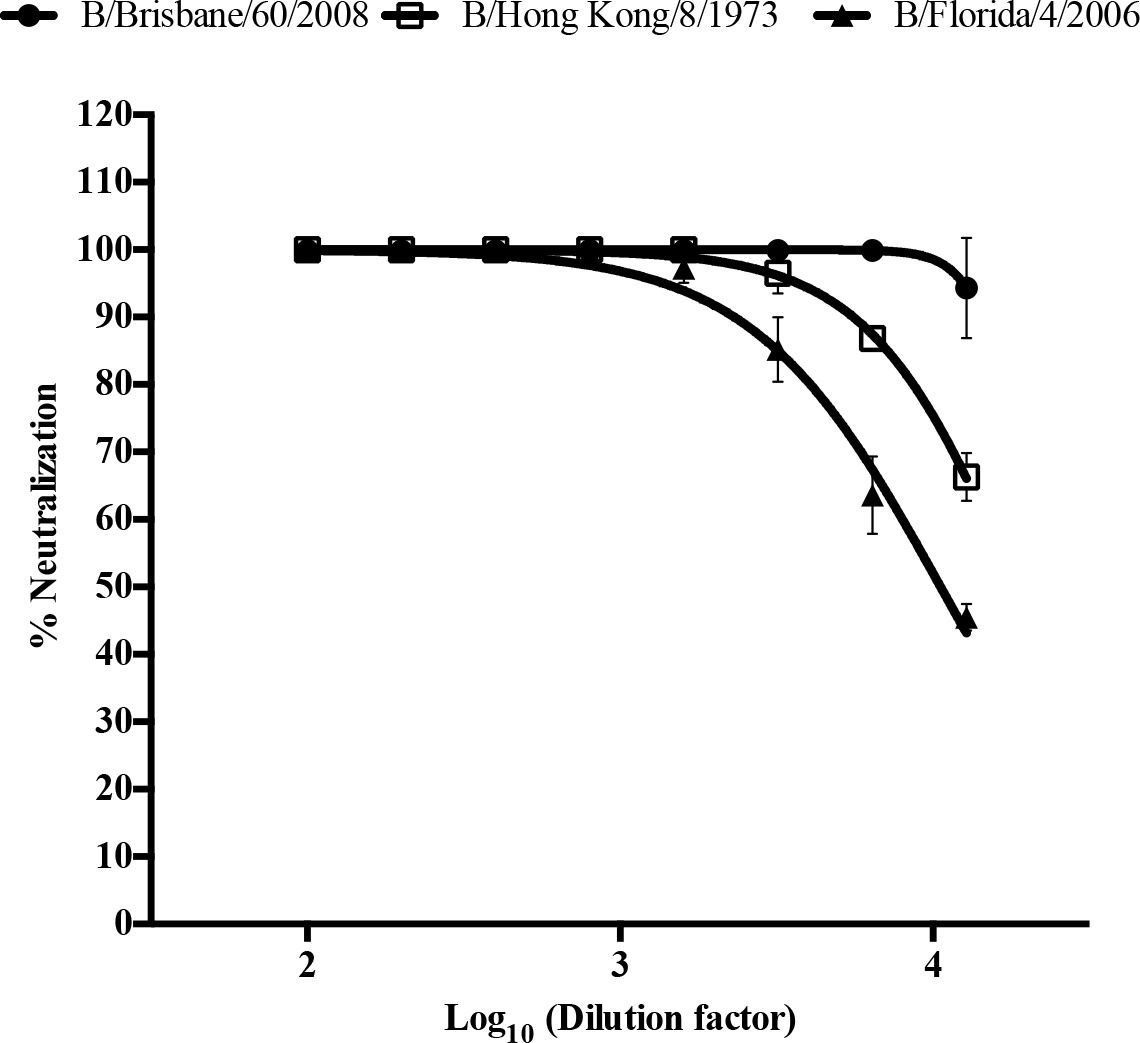
Anti-B/Brisbane/60/2008 reference serum neutralizes different influenza B PV with differing specificity. B/Brisbane/60/2008 (Victoria lineage) HA PV were neutralized strongly, followed by B/Hong Kong/8/1973 (Pre lineage split) and B/Florida/4/2006 (Yamagata lineage).

Specificity and sensitivity are further confirmed by the testing of lineage specific anti-influenza B monoclonal antibodies (Verma 2017). These mAbs were tested on all strains of influenza B mentioned previously, with very potent neutralisation recorded against each targeted lineage respectively. mAb 2F11 which binds to an epitope conserved between both lineages, neutralises all PVs with the exception of A/Brisbane/60/2008. With the exception of A/Brisbane/60/2008 which is not neutralised by the Victoria specific mAbs (8E12, 5A1), B/Victoria/2/1987 and the pre-lineage strain B/Hong Kong/8/1973 are neutralized. Yamagata specific mAbs potently neutralise Yamagata lineage PV B/Yamagata/16/1988, B/Bangladesh/3333/2007 and B/Florida/4/2006. See Figure 7. These results suggest that the Victoria lineage has maintained the B/Hong Kong/8/1973 epitopes targeted by mAbs 8E12 and 5A1. These results confirm that the pMN assay is detecting crossreactive neutralising antibodies when testing serum, as single-epitope neutralisation assays show no cross-reactivity whatsoever. Further work is required to evaluate the unexpected results generated with the Victoria lineage strain B/Brisbane/60/2008.

**Figure 7.**
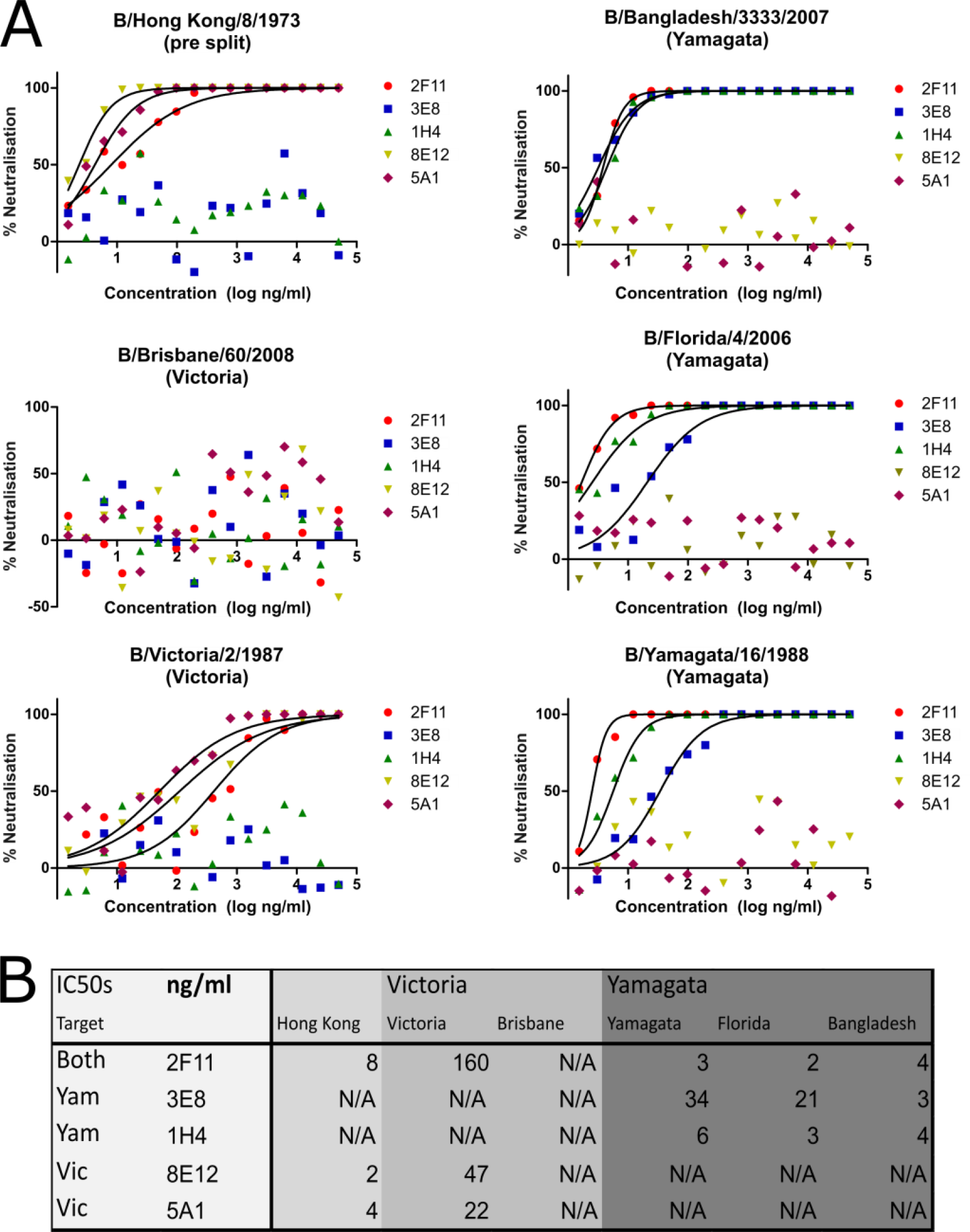
Lineage specific monoclonal antibodies neutralise influenza B pseudotypes with high specificity. **A)** Yamagata specific mAbs neutralize B/Bangladesh/333/2007, B/Florida/4/2006 and B/Yamagata/16/1988 whereas Victoria specific mAbs neutralize B/Victoria/2/1987 as well as the prelineage split strain B/Hong Kong/8/1973. B/Brisbane/60/2009 (Victoria) is not neutralized by any of the listed mAbs. **B)** IC_50_ values (ng/ml) calculated using nonlinear regression based on data shown in A.

### Influenza B pMN does not correlate with haemagglutination inhibition assay

Human sera from NCT00942071 clinical trial were screened against the vaccine-matching B/Brisbane/60/2008 in a pMN assay. Log10 (IC_50_) and IC_50_ values were calculated and log10 (IC_50_) values were then compared with the log10 HI titres (Antrobus et al. 2013). Surprisingly, high neutralization responses were detected against the vaccination strain B/Brisbane/60/2007 in both the post-vaccination and pre-vaccination sera. Furthermore, analysing the correlation between the HI and pMN assays, discordant correlation (r = 0.1632, p = 0.3563) was observed (Figure 8).

**Figure 8.**
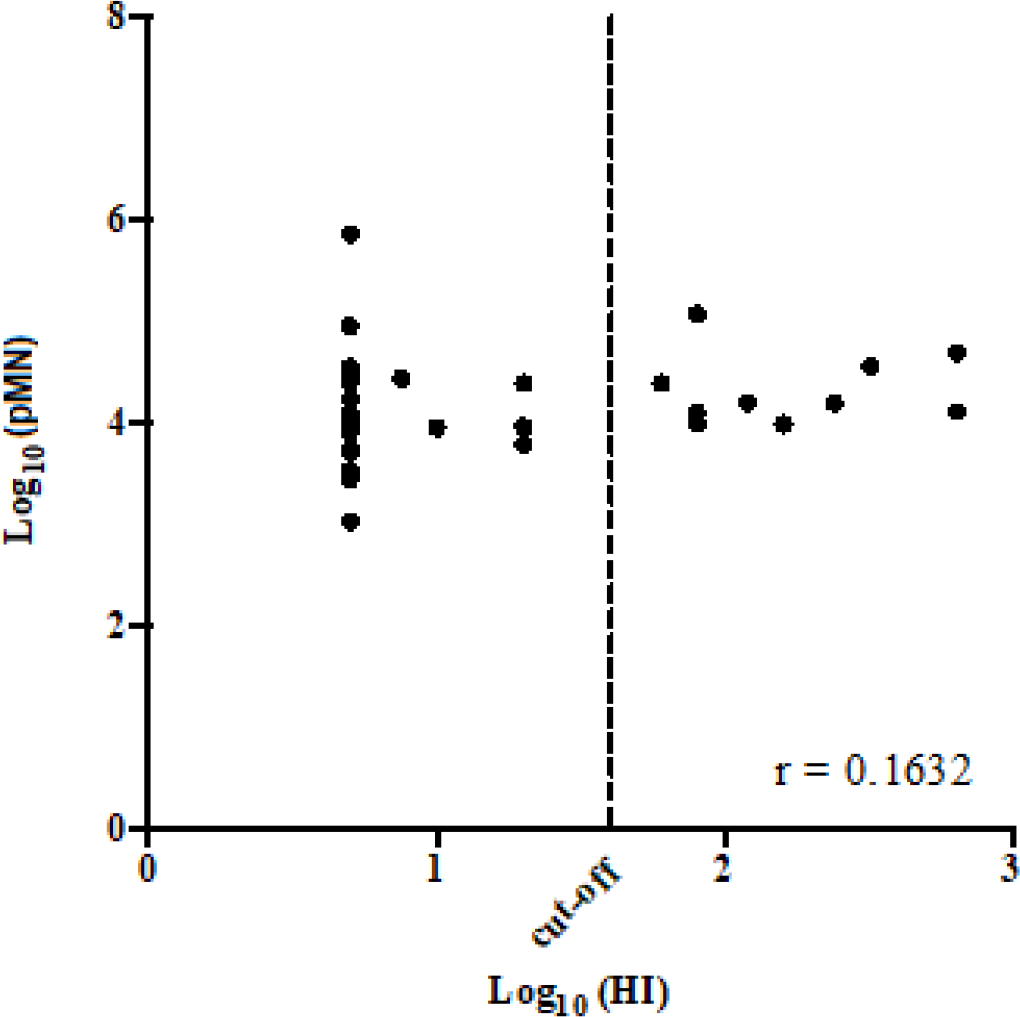
Correlation of HI and pMN assays performed with the B/Brisbane/60/2008 influenza strain. The cut off reported on the log10 (HI) axis corresponds to log10 (40).

The discordant correlation observed is not surprising: the influenza B HI assay has previously been demonstrated to be insensitive (Wood et al. 1994; Mancini et al. 1983; Oxford et al. 1982), whereas pMN is highly sensitive (Garcia et al. 2010; Wang et al. 2008; Temperton et al. 2007). The HI insensitivity is also demonstrated by the fact that it was unable to detect antigenic differences between influenza B viruses until the 1980s (Rota et al. 1990). Meanwhile, phylogenetic analyses have shown that the two distinct lineages were already present in the second half of the 1970s (Chen & Holmes 2008). SRH and classical HI were shown previously to correlate poorly with influenza B (Oxford et al. 1982). More recently, it was reported that discordant correlation could be observed between SRH, MN, HI, and pMN, since all these assays measure different antibody responses (Molesti et al. 2014). It will be necessary to test other sets of sera to understand the correlation of the Influenza B pMN with HI. Since HI is able to detect only head-directed antibodies, whereas pMN is able to detect antibody against the full HA, it will also be necessary to evaluate if the pMN assay correlates with SRH or MN. Of special interest would be the correlation study with MN since pMN and MN are based on the same principle, differing essentially in sensitivity, and are both able to detect stalk-directed cross-neutralizing antibodies (Corti et al. 2011).

### Influenza B pMN detects cross-reactive antibody responses between Victoria and Yamagata lineages

The data obtained (Figure 9A) in pMN assays using B/Brisbane/60/2008 show that strong neutralizing antibody responses are present pre-vaccination and that overall, the vaccination itself fails (p = 0.5791) to induce higher antibody responses. However a shift of the IC_50_ distribution first quartile can be observed post-vaccination. Furthermore, when the samples were stratified for the vaccine regimens administered to the subjects (Figure 9D) no statistical significance was detected between the TIV + placebo and TIV + MVA-NP+M1 groups, as already observed during the previous analysis of HI data (Antrobus et al. 2013). Different studies have shown the influenza A pMN is more sensitive at detection of cross-reactive and stalk-directed antibody responses than traditional serological assays. For this reason, influenza B pMN was evaluated in the same manner, by using sera from the NCT00942071 clinical trial. This took the form of neutralization assays with influenza B viruses not included in the trivalent vaccine that was originally administered to the test subjects.

**Figure 9:**
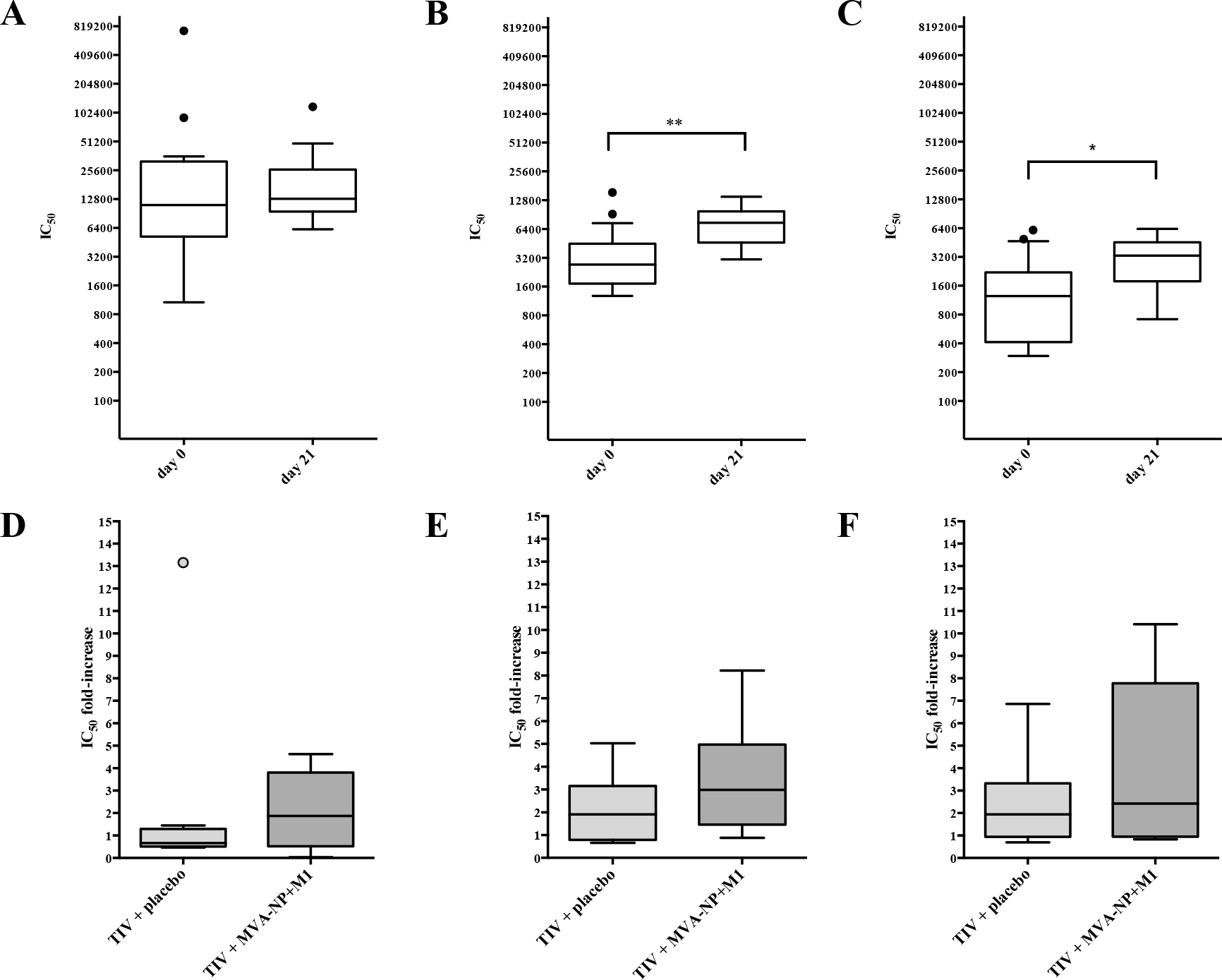
Evaluation of NCT00942071 clinical trial sera with influenza B pMN assay. IC_50_ and measured pre- (day 0) and post- (day 21) vaccination using as antigen B/Brisbane/60/2008 PV (A), B/Hong Kong/8/1973 PV (B), and B/Florida/4/2006 PV (C) were compared using Box-and-Whisker plots. After data stratification on tha basis of vaccine regimens, the IC_50_ fold-increases for B/Brisbane/60/2008 PV (D), B/Hong Kong/8/1973 PV (E), and B/Florida/4/2006 PV (F) of each group were also compared

NCT00942071 samples were evaluated against B/Hong Kong/8/1973 PVs, that has antigenic characteristics typical of influenza B strains circulating before the lineage split. Samples were also tested against a strain of the Yamagata lineage (B/Florida/4/2006). For B/Hong Kong/8/1973, antibody responses were detected at vaccination day 0 and at day 21 with post-vaccination neutralizing titres exhibiting a significant increase (p = 0.0046) (Figure 9B). For B/Florida/4/2006 the neutralizing responses observed at day 0 increased significantly (p = 0.0129) at day 21 (Figure 9C). After further analysis of the data obtained, no statistical differences were observed between the two vaccine regimens used (TIV + placebo and TIV + MVA-NP+M1) as observed in the B/Brisbane/60/2008 pMN (Figure 9E-F). With regard to the age of the participants in this clinical trial (50 years and above) (Antrobus et al. 2013), it is likely that the subjects have encountered the two strains (B/Florida/4/2006 and B/Hong Kong/8/1973) previously, so it is to be expected that a certain level of antibody against theses two strains could already be present. However, the high titres observed, especially against the B/Hong Kong/8/1973 PVs, and the increase in the neutralization titre after vaccination, indicate that a certain level of cross-reactivity could be present.

Before the Yamagata and Victoria lineage split, SRH was shown to be able to detect seroconversion, additionally for strains not encountered previously, and not included in the administered vaccines (Oxford et al. 1982). In that case, this was not interpreted as lower specificity of the assay, but as a characteristic of influenza B virus to be able to induce a cross-reactive response (Oxford et al. 1982). Recently it was shown that seroconversion and concomitant increase in antibody response was observed against a Yamagata strain. This occurred when a trivalent vaccine containing a Victoria lineage strain was administered after priming with a Yamagata-strain containing vaccine (Skowronski et al. 2012). Here, analogously, when a Victoria-strain based vaccine (B/Brisbane/60/2008) is administered, a significant increase in the neutralizing antibody response against a Yamagata strain (B/Florida/4/2006) and a pre-lineage division strain (B/Hong Kong/8/1973) is observed, whereas the increase against the vaccine strain, if present, is not statistically significant. This could indicate that the cross-reactivity between influenza B viruses is primarily related to an “antigenic-sin”. However, it also indicates that influenza B responses could be driven by antibodies able to cross-react between different influenza B strains.

### A panel of monoclonal antibodies discriminate between influenza B lineages using pMN

At present, monoclonal antibodies able to cross-neutralize the two influenza B lineages have been isolated from humans and it was demonstrated that they recognise epitopes present at the hinge of the HA head and stalk region or in the stalk region itself (Dreyfus et al. 2013). However, serological studies have failed to identify these influenza B antibodies mainly as a result of the insensitivity of the classical experimental tools for stalk antibody detection. On the contrary, pMN is highly sensitive in detecting stalk antibodies and the results observed could potentially highlight the presence of such antibodies in human sera. Further studies should therefore be conducted to verify the real spread of these antibodies and to confirm the results observed. These studies could be enhanced by using an influenza B version of an influenza A chimeric PV assay that can delineate functional antibody responses directed against the head from those directed against the stalk (Ferrara & Temperton 2017).

### Evaluation of a panel of Caspian Sea seal sera for neutralizing responses against influenza B PVs

Seal sera obtained from the Caspian Sea were evaluated for neutralizing activity against influenza B strains (B/HongKong/8/1973, B/Yamagata/16/1988, B/Victoria/2/1987) using pMN (Table 2). Unlike the situation in the human vaccine sera (NCT00942071), where pre-vaccination titres were already high, this was not the case in the limited number of seal sera tested. For these, only a single positive signal was observed, against B/Victoria (IC_50_ = 44). Further studies will be necessary; including serology undertaken using pMN against HAs from strains circulating within seal populations from the Caspian Sea area.

## CONCLUSION

The data presented in this paper demonstrate the successful production of influenza B PVs, which are produced at high titre using co-transfected proteases in a similar fashion to influenza A PVs. These useful tools can be used to measure “clean” immune responses against the HA in the absence of the native NA, with strain and lineage specificity as well as cross-reactive antibody responses detected and distinguishable. However, more experiments are necessary to understand the advantages and disadvantages of the influenza B pMN assay in comparison with classical serological methods.

## ACKNOWLEDGEMENTS

The authors thank Jerry Weir, U.S. Food and Drug Administration, 10903 New Hampshire Avenue, Silver Spring, MD 20993, United States of America, for the kind provision of influenza B mAbs described in Verma 2017.

